# PIVOT: Platform for Interactive Analysis and Visualization of Transcriptomics Data

**DOI:** 10.1101/053348

**Authors:** Qin Zhu, Stephen A Fisher, Hannah Dueck, Sarah Middleton, Mugdha Khaladkar, Junhyong Kim

**Affiliations:** Department of Biology, University of Pennsylvania, 304G Lynch Laboratory, 433 S University Avenue, Philadelphia, PA, USA.

**Keywords:** Transcriptomics, graphical user interface, interactive visualization, exploratory data analysis

## Abstract

Many R packages have been developed for transcriptome analysis but their use often requires familiarity with R and integrating results of different packages is difficult. Here we present PIVOT, an R-based application with a uniform user interface and graphical data management that allows non-programmers to conveniently access various bioinformatics tools and interactively explore transcriptomics data. PIVOT supports many popular open source packages for transcriptome analysis and provides an extensive set of tools for statistical data manipulations. A graph-based visual interface is used to represent the links between derived datasets, allowing easy tracking of data versions. PIVOT further supports automatic report generation, publication-quality plots, and program/data state saving, such that all analysis can be saved, shared and reproduced.

## Introduction

Technologies such as RNA-sequencing and microarray measure gene expressions and present them as high-dimensional expression matrixes for downstream analyses. In recent years, many programs have been developed for the statistical analysis of transcriptomics data, such as edgeR [1] and DESeq [2] for differential expression testing, and monocle [3], Seurat [4] and SCDE [5] for single cell RNA-Seq data analysis. Besides these, the Comprehensive R Archive Network (CRAN) [6] and Bioconductor [7] host various statistical packages addressing different aspects of transcriptomics study and provides recipes for a multitude of analysis workflows. Making use of these R analysis packages requires expertise in R and often custom scripts to integrate the results of different packages. In addition, many exploratory analyses of transcriptome data involve repeated data manipulations such as transformations (e.g., normalizations), filtering, merging, etc., each step generating a derived dataset whose version and provenance must be tracked. Previous efforts to address these problems include designing standardized workflows [8], building a comprehensive package [4] or assembling pipelines into integrative platforms such as Galaxy [9] or Illumina BaseSpace [10]. Designing workflows or using large packages still requires a significant amount of programming skills and it can be difficult to make various components compatible or applicable to specific datasets. Integrative platforms offer greater usability but trades off flexibility, functionality and efficiency due to limitations on data size, parameter choice and computing power. For example, the Galaxy platform is designed as discrete functional modules which require separate file inputs for different analysis. This design not only makes user-end file format conversion complicated and time-consuming, but also breaks the integrity of the analysis workflow, limiting the sharing of global parameters, filtering criteria and analysis results between modules. Tools such as ASAP [11] and DEApp [12] provides an interactive graphical interface for a small number of packages. But, these and other similar packages all adopt a rigid workflow design and have limited data provenance tracking, and none of the packages provide mechanisms for saving, sharing and reproducing analysis results. Furthermore, many web-based applications require users to upload data to a server, which might be prohibited by HIPPA (Health Insurance Portability and Accountability Act of 1996) for clinical data analysis.

Here we developed PIVOT, an R-based platform for exploratory transcriptome data analysis. We leverage the Shiny framework [13] to bridge open source R packages and JavaScript-based web applications, and to design a user-friendly graphical interface that is consistent across statistical packages. The Shiny framework translates user-driven events (e.g. pressing buttons) into R interpretable reactive data objects, and present results as dynamic web content. PIVOT incorporates four key features that assists user interactions, integrative analysis and provenance management:

- PIVOT directly integrates existing open source packages by wrapping the packages with a uniform user-interface and visual output displays. The user interface replaces command line options of many packages with menus, sliders, and other option controls, while the visual outputs provide extra interactive features such as change of view, active objects, and other user selectable tools.
- PIVOT provides many tools to manipulate a dataset to derive new datasets including different ways to normalize a dataset, subset a dataset, etc. In particular, PIVOT supports manipulating the datasets using the results of an analysis; for example, an user might use the results of differential gene expression analysis to select all gene satisfying some p-value filter. PIVOT implements a visual data management system, which allows users to create multiple data views and graphically display the linked relationship between data variants, allowing navigation through derived data objects and automated re-analysis.
- PIVOT dynamically bridges analysis modules to allow results from one module being used as inputs for another. Thus, it provides a flexible framework for users to combine tools into customizable pipelines for various analysis purposes.
- PIVOT provides facilities to automatically generate reports, publication-quality figures, and reproducible computations. All analyses and data generated in an interactive session can be packaged as a single R object that can be shared to exactly reproduce any results.

## Methods

### Data Input and Transformations

Read counts obtained from RNA-Seq quantification tools such as HTSeq [14] or featureCounts [15] can be directly uploaded into PIVOT as text, csv or Excel files. PIVOT automatically merges sample counts into an expression matrix and performs user selected data transformations including normalization, log transformation, or standardization. We have included multiple RNA-Seq data normalization methods including DESeq normalization [16], trimmed mean of M-values (TMM) [17], quantile normalization [18], RPKM/TPM [19], Census normalization [20], and Remove Unwanted Variation (RUVg) [21] (Table 1). If samples contain spike-in control mixes such as ERCC [22], PIVOT will also separately analyze the ERCC count distribution and allow users to normalize the data using the ERCC control. Existing methods can be customized by the user by setting detailed normalization parameters. For example, we implement a modification of the DESeq method by making the inclusion criterion a user set parameter, making it more applicable to sparse expression matrices such as single cell RNA-Seq data [23].

**Table 1.** List of tools currently integrated in PIVOT.

| **PIVOT Modules** | **Tools Integrated** |
| --- | --- |
| **Normalization** | DESeq, Modified DESeq, TMM, Upper quartile, CPM/RPKM/TPM, RUV, Spike-in regression, Census |
| **Feature/Sample Filtering** | List based, Expression based and Quality based filters |
| **Basic Analysis Modules** | Data distribution plots, Dispersion Analysis, Rank-frequency plot, Spike-in Analysis, Feature heatmap, etc. |
| **Differential Expression** | DESeq2, edgeR, SCDE, Monocle, Mann-Whitney U test |
| **Clustering/Classification** | Hierarchical, K-means, Community detection, Classification with caret, Cell state ordering with Monocle2 |
| **Dimension Reduction** | PCA, t-SNE, Metric/Non-Metric MDS, penalized LDA |
| **Correlation Analysis** | Pairwise scatter plots, Sample/feature correlation heatmap, Co-expression analysis |
| **Gene Set Enrichment Analysis** | KEGG pathway analysis, Gene ontology analysis |
| **Network Analysis** | STRING protein association network, Regnetwork visualization, Mogrify based trans-differentiation factor prediction |
| **Toolkit** | BioMart gene annotation query, Venn diagram |

Users can upload experiment design information such as conditions and batches, which can be visualized as annotation attributes (e.g., color points/sidebars) or used as model specification variables for downstream analyses such as differential expression. PIVOT supports flexible operations to filter data for row and column subsets as well as for merging datasets, creating new derived datasets. Multiple summary statistics are automatically computed to help users identify possible outliers. Users can manually select samples for analysis, or specify statistical criteria on analysis results such as expression threshold, dispersion cutoff, Cook’s distance or size factor range to remove unwanted features and samples.

### Visual Data Management with Data Map

When analyzing large datasets, a common procedure is to first perform quality control to remove low quality elements, then normalize the data and finally generate different data subsets for various analysis purposes. Some analyses require filtering out genes with low expressions, while others are designed to be performed on a subset of the genes such as transcription factors. During secondary analyses, outliers may be detected requiring additional scrutiny. All these data manipulations generate a network of derived datasets from the original data and require a significant amount of effort to track. Failure to track the data lineage could affect the reproducibility and reliability of the study. Furthermore, an investigator might wish to repeat an analysis over a variety of derived datasets, which may be tedious and error-prone to carry out manually. To address this problem, we implemented a graphical data management system in PIVOT.

As the user generates derived datasets with various data manipulations, PIVOT records and presents the data provenance in an interactive tree graph, the “Data Map”. As shown in Figure 1, each node in the data map represents a derived dataset and the edges contain information about the details of the derivation operation. Users can attach analysis results to the data nodes as interactive R markdown reports [24] and switch between different datasets or retrieve analysis reports by simply clicking the nodes. Upon switch to a new dataset selected from the Data Map, PIVOT automatically re-runs analyses and updates parameter choices when needed. Thus, a user can easily compare results of a workflow across derived datasets. The data map is generated with the visNetwork package [25] and can be directly edited, so that users can rename nodes, add notes, or delete data subsets and analysis reports that are no longer useful. The full data history is also presented as downloadable tables with all sample and feature information as well as data manipulation details.

**Fig. 1.**
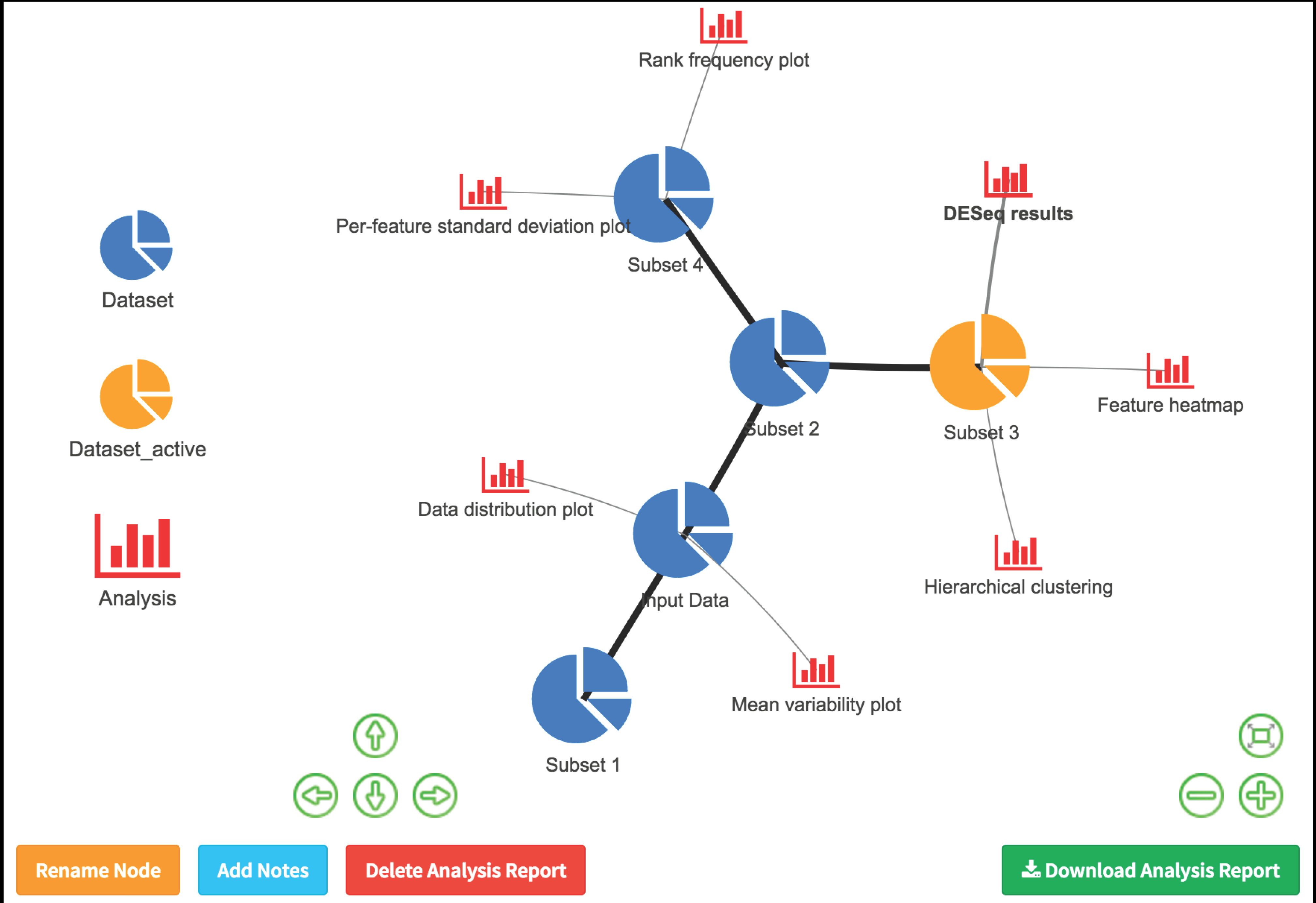
Data management with data map. The map shows the history of the data change and the association between analysis and data nodes. Users can hover over edges to see operation details, or click nodes to get analysis reports or switch active subsets.

### Comprehensive Toolset for Exploratory Analysis

PIVOT is designed to aid exploratory analysis for both single cell and bulk RNA-Seq data, thus we have incorporated a large set of commonly used tools (see Table 1). PIVOT supports many visual data analytics including QC plots (number of detected genes, total read counts and estimated size factors; Fig. 2a, data from [26]), transcriptome statistics plots (e.g., rank-frequency plots, mean-variability plots, etc; Fig. 2b), and sample and feature correlation plots (e.g., heatmaps, smoothened scatter plots, etc.). All visual plots feature interactive options and a query function is provided which allows users to search for features sharing similar expression patterns with a target feature. PIVOT provides users extensive control over parameter choices. Each analysis module contains multiple visual controls allowing users to adjust parameters and obtain updated results on the fly.

**Fig. 2.**
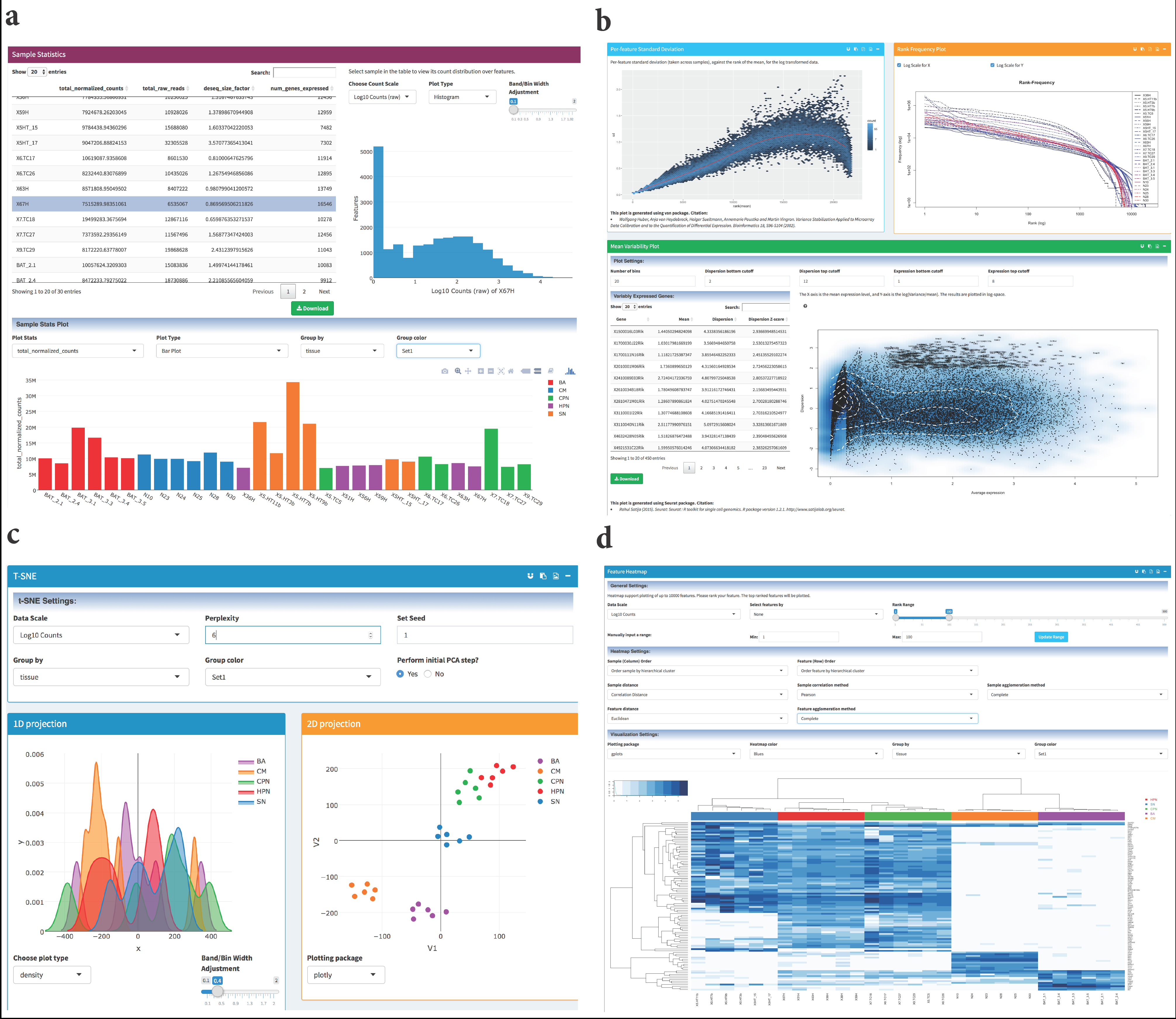
Selected analysis modules in PIVOT. (a) The table on the left lists basic sample statistics. The selected statistics are plotted below the table, and clicking a sample in the table will plot its count distribution. (b) Mean-Standard deviation plot (top left, with vsn package), rank frequency plot (top right) and mean variability plot (bottom, with Seurat package). (c) The t-SNE module plots 1D, 2D and 3D projections (3D not shown due to space). (d) Feature heatmap with the top 100 differentially expressed genes reported by DESeq2 likelihood ratio test.

### Integrative Analysis and Interactive Visualization

PIVOT transparently bridges multiple sequences of analyses to form customizable analysis pipelines. For example, with single cell data collected from heterogeneous tumor or tissue, a user can first perform PCA or t-SNE [27] (Fig. 2c) to visualize the low dimensional embedding of the data. If there is clear clustering pattern, possibly originated from different cell types, the user can directly specify cell clusters by dragging selection boxes on the graph, or perform K-means or hierarchical clustering with the projection matrix. One can proceed to run DE or penalized LDA [28] to identify cluster-specific marker genes, which can then be used to filter the datasets for generating a heatmap showing distinctive expression pattern across cell types (Fig. 2d). Within each determined cell type, a user may further apply the walk-trap community detection method [29] to identify densely connected network of cells, which are indicative of potential subpopulations [30].

As another example, for time-series data such as cells collected at different stages of development or differentiation, one can use monocle [3], which implements an unsupervised algorithm for pseudo-temporal ordering of single cells [31]. We have incorporated the latest monocle 2 workflow in PIVOT, including cell state ordering, unsupervised cell clustering, gene clustering by pseudo-temporal expression pattern and cell trajectory analysis. Besides the DE method implemented in monocle, one can also run DESeq, edgeR, SCDE or the Mann-Whitney U test. A user can specify whether to perform basic DE analysis or a multi-factorial DE analysis with customized formulae for complex experimental designs such as time-series or controlling for batch effects. Results are presented as dynamic tables including all essential statistics such as maximum likelihood estimation and confidence intervals. Each gene entry in the table can be clicked and visualized as violin plots or box plots, showing the actual expression level across conditions. Once DE results are obtained, the user can further explore the connections between DE genes and identify potential trans-differentiation factors as introduced in the Mogrify algorithm [32]. PIVOT provides several extensions of functionality from the original Mogrify method. The network analysis module allows users to plot the log fold changes (LFC) of DE genes in a protein-protein interaction network obtained from the STRING database (Fig. 3a) [33] or a directed regulatory network graph constructed from the Regnetwork repository (Fig. 3b) [34]. With scoring based on the p-value and log fold change, the graph can be zoomed to only include top-rank genes, showing the regulatory “hot spot” of the network. PIVOT provides users with multiple options for defining the network influence score of transcription factors, and will produce lists of potential trans-differentiation factors based on the final ranking. As shown in Fig. 3c, with the FANTOM5 expression data of fibroblasts and ES cells [35], PIVOT correctly reports OCT4 (POU5F1), NANOG and SOX2 as key factors for trans-differentiation [36]. In addition to the DESeq results used by the original Mogrify algorithm, a user can choose to use SCDE or edgeR results to perform trans-differentiation analysis on single cell datasets.

**Fig. 3.**
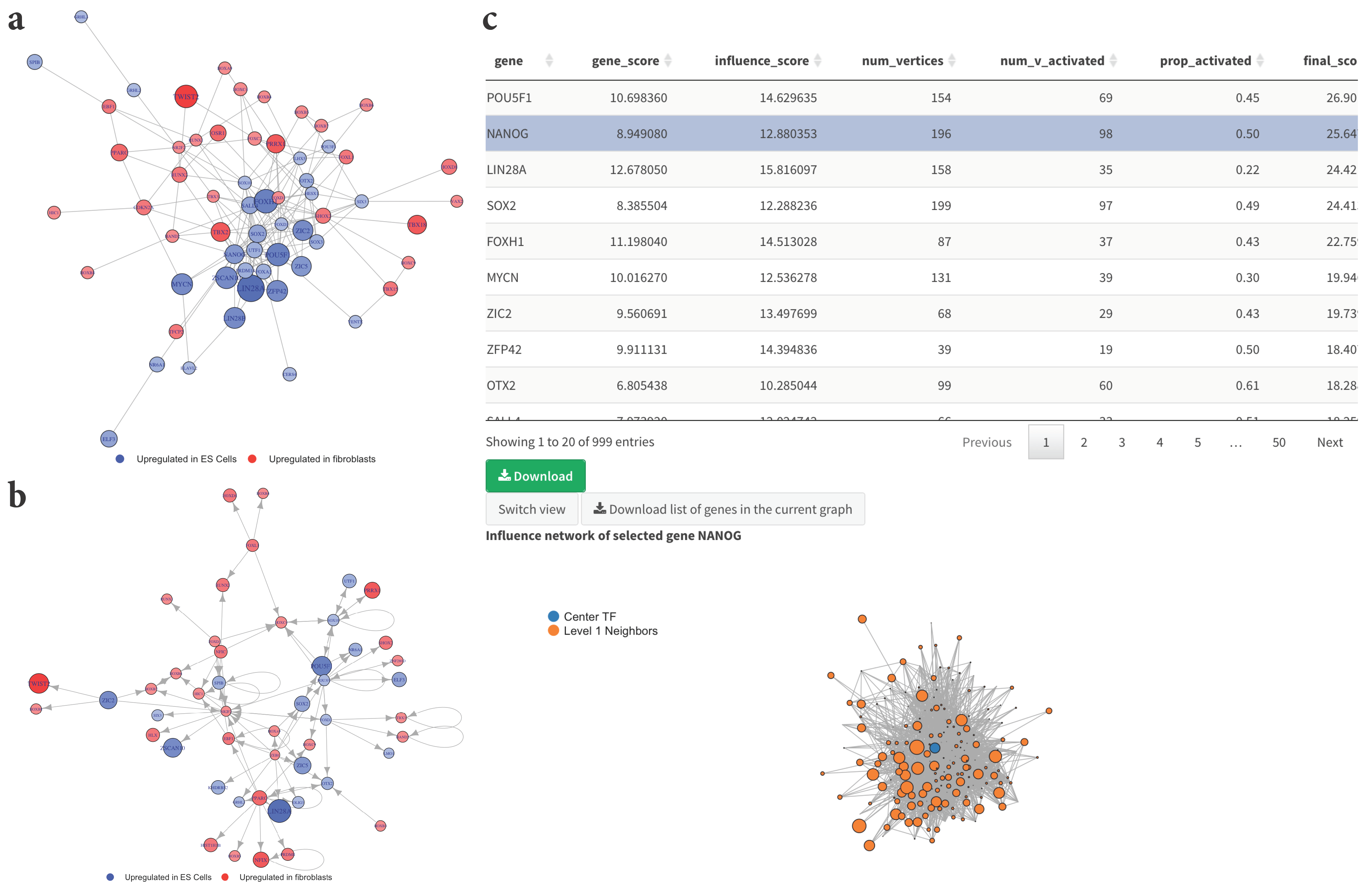
Network analysis for the identification of potential transdifferentiation factors. (a, b)Graphs showing the connection between transcription factors differentially expressed between fibroblasts and ES cells. 3a is an undirected graph showing the protein-protein interaction relationship based on the STRING database, and 3b is constructed based on the Regnetwork repository, showing the regulatory relationship. The size of the nodes and the color gradient indicate the log fold change of the genes. The graphs have been zoomed in to only include the genes with large LFC and small p-value. (c) Predicted transdifferentiation factor lists based on the network score ranking. The table includes information such as the center transcription factor score, the total number of vertices in its direct neighborhood, and the number of activated neighbors with gene score above a user-specified threshold. Clicking entries on the table will plot the local neighborhood network centered on that TF.

Another useful feature of PIVOT is that it provides users multiple visualization options by exploiting the power of various plotting packages. For example, users can either generate publication-quality heatmap graphs (implemented in gplots package [37]), or interactively explore the heatmap with the heatmaply view [38]. For principal component analysis, PIVOT uses three different packages to present the 2D and 3D projections. The plotly package [39] displays sample names and relevant information as mouse-over labels, while the ggbiplot [40] presents the loadings of each gene on the graph as vectors. The threejs package [41] fully utilizes the power of WebGL and outputs rotatable 3D projections. In the network analysis module, we utilize both igraph [42] and networkD3 [43] package to plot the transcription factor centered local network. The latter provides a force directed layout, which allows users to drag the nodes and visualize the physical simulation of the network response.

### Reproducible Research and Complete Provenance Capture

PIVOT automatically records all data manipulations and analysis steps. Once an analysis has been performed, users will have the option of pasting related R markdown code to a shinyAce report editor [44], or download the report as either a pdf or interactive html document. All results and associated parameters will be captured and saved to the report along with user-provided comments. PIVOT states are automatically saved in cases of browser refresh, crash or user exit, and can also be manually exported, shared and loaded. Thus, all analyses performed in PIVOT are fully encapsulated and can be shared or disseminated as a single data+provenance object, allowing universally reproducible research.

### Implementation

PIVOT is written in R and is distributed as an R package. It is developed using the Shiny framework, multiple R packages and a collection of scripts written by members of J. Kim’s Lab at University of Pennsylvania. PIVOT exports multiple Shiny modules [45] which can be used as design blocks for other Shiny apps, as well as R functions for transcriptomics analysis and plotting. A proficient R user can easily access data objects, analysis parameters and results exported by PIVOT and use them in customized scripts. PIVOT has been tested on macOS, Linux and Windows. It can be downloaded from Kim Lab Software Repository (http://kim.bio.upenn.edu/software) or GitHub (https://github.com/qinzhu/PIVOT).

### Discussion and conclusions

We developed PIVOT for easy, fast, and exploratory analysis of the transcriptomics data.

Toward this goal we have automated the analysis procedures and data management, and we provide users with detailed explanations both in tooltips and a user manual. PIVOT exploits the power of multiple plotting packages and gives users full control of key analysis and plotting parameters. Given user input that leads to function errors, PIVOT will alert the user and provide corrective suggestions. PIVOT states and reports can be shared between researchers to facilitate the discussion of expression analysis and future experimental design. PIVOT is designed to be extensible and future versions will continue to integrate popular transcriptome analysis routines as they are made available to the research community.

## Abbreviations

PIVOT: Platform for Interactive analysis and Visualization Of Transcriptomics data
TMM: trimmed mean of M-values
RPKM: Reads Per Kilobase per Million mapped reads
RUV: remove unwanted variation
DE: differential expression
GUI: graphical user interfaces
PCA: principal component analysis
t-SNE: t-Distributed Stochastic Neighbor Embedding
LDA: linear discriminant analysis
MDS: multidimensional scaling
ES cells: Embryonic stem cells
TF: transcription factor
LFC: log fold change

## Declarations

### Acknowledgements

We are grateful to all members in Junhyong Kim’s lab and James Eberwine’s lab for their participation in the beta-testing of the program and their valuable feedbacks and suggestions. This research has been supported by NIMH U01MH098953 grant to J. Kim and J. Eberwine.

### Availability of data and materials

Project name: PIVOT

Project home page: http://kim.bio.upenn.edu/software/pivot.shtml Operating systems: macOS, Linux, Windows.

Programming language: R

Other requirements: Dependent R packages.

License: GNU.

### Authors’ contributions

QZ carried out the programming tasks. QZ, SF and JK designed the application. SF, HD, SM and MK contributed scripts and extensive software testing. QZ, SF and JK wrote the manuscript.

## References

1. McCarthy DJ, Chen Y, Smyth GK. Differential expression analysis of multifactor RNA-Seq experiments with respect to biological variation. Nucleic Acids Research. 2012;40.

2. Love MI, Huber W, Anders S. Moderated estimation of fold change and dispersion for RNA-seq data with DESeq2. Genome biology. 2014;15:1–21.

3. Trapnell C, Cacchiarelli D, Grimsby J, Pokharel P, Li S, Morse M, Lennon NJ, Livak KJ, Mikkelsen TS, Rinn JL. The dynamics and regulators of cell fate decisions are revealed by pseudotemporal ordering of single cells. Nature biotechnology. 2014;32:381–386.

4. Satija R, Farrell JA, Gennert D, Schier AF, Regev A. Spatial reconstruction of single-cell gene expression data. Nature biotechnology. 2015;33:495–502.

5. Kharchenko PV, Silberstein L, Scadden DT. Bayesian approach to single-cell differential expression analysis. Nature methods. 2014;11:740–742.

6. Hornik K. The comprehensive R archive network. Wiley Interdisciplinary Reviews: Computational Statistics. 2012;4:394–398.

7. Huber W, Carey VJ, Gentleman R, Anders S, Carlson M, Carvalho BS, Bravo HC, Davis S, Gatto L, Girke T. Orchestrating high-throughput genomic analysis with Bioconductor. Nature methods. 2015;12:115–121.

8. Anders S, McCarthy DJ, Chen Y, Okoniewski M, Smyth GK, Huber W, Robinson MD. Count-based differential expression analysis of RNA sequencing data using R and Bioconductor. Nature protocols. 2013;8:1765–1786.

9. Goecks J, Nekrutenko A, Taylor J. Galaxy: a comprehensive approach for supporting accessible, reproducible, and transparent computational research in the life sciences. Genome Biology. 2010;11:R86.

10. Illumina basespace. https://basespace.illumina.com/home/index. Accessed 8 June 2017.

11. Gardeux V, David FP, Shajkofci A, Schwalie PC, Deplancke B. ASAP: a Web-based platform for the analysis and interactive visualization of single-cell RNA-seq data. Bioinformatics. 2017:btx337.

12. Li Y, Andrade J. DEApp: an interactive web interface for differential expression analysis of next generation sequence data. Source code for biology and medicine. 2017;12:2.

13. Chang W, Cheng J, Allaire J, Xie Y, McPherson J. shiny: Web Application Framework for R. CRAN. 2017.

14. Anders S, Pyl PT, Huber W. HTSeq–A Python framework to work with high-throughput sequencing data. Bioinformatics. 2014:btu638.

15. Liao Y, Smyth GK, Shi W. featureCounts: an efficient general purpose program for assigning sequence reads to genomic features. Bioinformatics. 2014;30:923–930.

16. Anders S, Huber W. Differential expression analysis for sequence count data. Genome Biology. 2010;11.

17. Robinson MD, Oshlack A. A scaling normalization method for differential expression analysis of RNA-seq data. Genome biology. 2010;11:1.

18. Dillies M-A, Rau A, Aubert J, Hennequet-Antier C, Jeanmougin M, Servant N, Keime C, Marot G, Castel D, Estelle J. A comprehensive evaluation of normalization methods for Illumina high-throughput RNA sequencing data analysis. Briefings in bioinformatics. 2013;14:671–683.

19. Mortazavi A, Williams BA, McCue K, Schaeffer L, Wold B. Mapping and quantifying mammalian transcriptomes by RNA-Seq. Nature methods. 2008;5:621–628.

20. Qiu X, Hill A, Packer J, Lin D, Ma Y-A, Trapnell C. Single-cell mRNA quantification and differential analysis with Census. Nature methods. 2017;14:309–315.

21. Risso D, Ngai J, Speed TP, Dudoit S. Normalization of RNA-seq data using factor analysis of control genes or samples. Nature biotechnology. 2014;32:896–902.

22. Lemire A, Lea K, Batten D, Gu JS, Whitley P, Bramlett K, Qu L. Development of ERCC RNA spike-in control mixes. Journal of biomolecular techniques: JBT. 2011;22.

23. Spaethling JM, Na Y-J, Lee J, Ulyanova AV, Baltuch GH, Bell TJ, Brem S, Chen HI, Dueck H, Fisher SA. Primary cell culture of live neurosurgically resected aged adult human brain cells and single cell transcriptomics. Cell reports. 2017;18:791–803.

24. Allaire J, Cheng J, Xie Y, McPherson J, Chang W, Allen J, Wickham H, Atkins A, Hyndman R. rmarkdown: Dynamic Documents for R. 2016.

25. Almende BV, Benoit T. visNetwork: Network Visualization using ‘vis.js’ Library. CRAN. 2016.

26. Dueck H, Khaladkar M, Kim TK, Spaethling JM, Francis C, Suresh S, Fisher SA, Seale P, Beck SG, Bartfai T. Deep sequencing reveals cell-type-specific patterns of single-cell transcriptome variation. Genome biology. 2015;16:1–17.

27. Maaten Lvd, Hinton G. Visualizing data using t-SNE. Journal of Machine Learning Research. 2008;9:2579–2605.

28. Witten DM, Tibshirani R. Penalized classification using Fisher’s linear discriminant. Journal of the Royal Statistical Society: Series B (Statistical Methodology). 2011;73:753–772.

29. Pons P, Latapy M. Computing communities in large networks using random walks. In International Symposium on Computer and Information Sciences. Springer; 2005:284–293.

30. Darmanis S, Sloan SA, Zhang Y, Enge M, Caneda C, Shuer LM, Gephart MGH, Barres BA, Quake SR. A survey of human brain transcriptome diversity at the single cell level. Proceedings of the National Academy of Sciences. 2015;112:7285–7290.

31. Magwene PM, Lizardi P, Kim J. Reconstructing the temporal ordering of biological samples using microarray data. Bioinformatics. 2003;19:842–850.

32. Rackham OJ, Firas J, Fang H, Oates ME, Holmes ML, Knaupp AS, Suzuki H, Nefzger CM, Daub CO, Shin JW. A predictive computational framework for direct reprogramming between human cell types. Nature genetics. 2016.

33. Szklarczyk D, Franceschini A, Wyder S, Forslund K, Heller D, Huerta-Cepas J, Simonovic M, Roth A, Santos A, Tsafou KP. STRING v10: protein–protein interaction networks, integrated over the tree of life. Nucleic acids research. 2014:gku1003.

34. Liu Z-P, Wu C, Miao H, Wu H. RegNetwork: an integrated database of transcriptional and post-transcriptional regulatory networks in human and mouse. Database. 2015;2015:bav095.

35. Lizio M, Harshbarger J, Shimoji H, Severin J, Kasukawa T, Sahin S, Abugessaisa I, Fukuda S, Hori F, Ishikawa-Kato S. Gateways to the FANTOM5 promoter level mammalian expression atlas. Genome biology. 2015;16:1.

36. Huangfu D, Osafune K, Maehr R, Guo W, Eijkelenboom A, Chen S, Muhlestein W, Melton DA. Induction of pluripotent stem cells from primary human fibroblasts with only Oct4 and Sox2. Nature biotechnology. 2008;26:1269–1275.

37. Warnes G, Bolker B, Bonebakker L, Gentleman R, Huber W, Liaw A, Lumley T, Maechler M, Magnusson A, Moeller S, et al. gplots: Various R Programming Tools for Plotting Data. CRAN. 2016.

38. Galili T. heatmaply: Interactive Heat Maps Using’plotly’. CRAN. 2017.

39. Sievert C, Parmer C, Hocking T, Chamberlain S, Ram K, Corvellec M, Despouy P. plotly: Create Interactive Web Graphics via ‘plotly.js’. CRAN. 2016.

40. Vu VQ. ggbiplot: A ggplot2 based biplot. R package. 2011.

41. Lewis BW. threejs: Interactive 3D Scatter Plots and Globes. CRAN. 2016.

42. Csardi G, Nepusz T. The igraph software package for complex network research. InterJournal, Complex Systems. 2006;1695:1–9.

43. Gandrud C, Allaire JJ, Russell K, Yetman C. networkD3: D3 JavaScript Network Graphs from R. CRAN. 2017.

44. LLC. TT. shinyAce: Ace editor bindings for Shiny. CRAN. 2016.

45. Cheng J. Modularizing Shiny app code. https://shinyrstudiocom/articles/moduleshtml. 2015.

